# Deamidation drives molecular aging of the SARS-CoV-2 spike receptor-binding motif

**DOI:** 10.1101/2021.05.20.445042

**Authors:** Ramiro Lorenzo, Lucas A. Defelipe, Lucio Aliperti, Stephan Niebling, Tânia F. Custódio, Christian Löw, Jennifer J. Schwarz, Kim Remans, Patricio O. Craig, Lisandro H. Otero, Sebastián Klinke, María García-Alai, Ignacio E. Sánchez, Leonardo G. Alonso

**Affiliations:** Centro de Investigación Veterinaria de Tandil (CIVETAN), CONICET-CICPBA-UNCPBA, Facultad de Ciencias Veterinarias, Universidad Nacional del Centro (FCV-UNCPBA), Tandil, Argentina; European Molecular Biology Laboratory, Hamburg Unit, 22607 Hamburg, Germany; Universidad de Buenos Aires, Consejo Nacional de Investigaciones Científicas y Técnicas, Instituto de Química Biológica de la Facultad de Ciencias Exactas y Naturales (IQUIBICEN), Facultad de Ciencias Exactas y Naturales, Laboratorio de Fisiología de Proteínas, Buenos Aires, Argentina; Centre for Structural Systems Biology, Notkestrasse 85, 22607 Hamburg, Germany; European Molecular Biology Laboratory, Heidelberg, Meyerhofstraße 1, 69117 Heidelberg, Germany; Universidad de Buenos Aires. Facultad de Ciencias Exactas y Naturales. Departamento de Química Biológica. Buenos Aires, Argentina; CONICET-Universidad de Buenos Aires. Instituto de Química Biológica de la Facultad de Ciencias Exactas y Naturales (IQUIBICEN). Buenos Aires, Argentina; Fundación Instituto Leloir, IIBBA–CONICET, and Plataforma Argentina de Biología Estructural y Metabolómica PLABEM, Av. Patricias Argentinas 435 (C1405BWE) Buenos Aires, Argentina; Instituto de Nanobiotecnologia (NANOBIOTEC), UBA-CONICET-Universidad de Buenos Aires, Buenos Aires, Argentina

## Abstract

The spike is the main protein component of the SARS-CoV-2 virion surface. The spike receptor binding motif mediates recognition of the hACE2 receptor, a critical infection step, and is the preferential target for spike-neutralizing antibodies. Post-translational modifications of the spike receptor binding motif can modulate viral infectivity and immune response. We studied the spike protein in search for asparagine deamidation, a spontaneous event that leads to the appearance of aspartic and isoaspartic residues, affecting both the protein backbone and its charge. We used computational prediction and biochemical experiments to identify five deamidation hotspots in the SARS-CoV-2 spike. Similar deamidation hotspots are frequently found at the spike receptor-binding motifs of related sarbecoviruses, at positions that are mutated in emerging variants and in escape mutants from neutralizing antibodies. Asparagine residues 481 and 501 from the receptor-binding motif deamidate with a half-time of 16.5 and 123 days at 37 °C, respectively. This process is significantly slowed down at 4 °C, pointing at a strong dependence of spike molecular aging on the environmental conditions. Deamidation of the spike receptor-binding motif decreases the equilibrium constant for binding to the hACE2 receptor more than 3.5-fold. A model for deamidation of the full SARS-CoV-2 virion illustrates that deamidation of the spike receptor-binding motif leads to the accumulation in the virion surface of a chemically diverse spike population in a timescale of days. Our findings provide a mechanism for molecular aging of the spike, with significant consequences for understanding virus infectivity and vaccine development.

## Introduction

In December 2019, a viral pneumonia outbreak was reported in Wuhan, China (1). This outbreak quickly turned into a pandemic disease (COVID-19) of international concerns (2), and the novel pathogen causative of a Severe Acute Respiratory Syndrome (SARS) was soon identified as SARS-CoV-2, a new member of the *Betacoronaviruses* genera. SARS-CoV-2 and SARS-CoV, the agent responsible for the 2002-2003 pneumonia outbreak, are closely related to the bat coronaviruses from which they likely originated and passed to an intermediate species that ultimately infected humans (3, 4). The host specificity and infectivity of SARS-CoV-2 and SARS-CoV rely on the spike protein (S).Through its receptor-binding domain (RBD, residues 319 to ~515), S recognizes the human Angiotensin-converting enzyme 2 (hACE2) with nanomolar affinity, triggering events that culminate with the fusion of the cellular and viral membranes (5). In SARS-CoV-2, the S protein is synthesized as a 1273-residue heavily glycosylated polypeptide that is cleaved by the host furin protease between the S1 (1-685) and S2 (686-1273) subunits (6). On the surface of native viruses, the S protein is mainly observed as a metastable trimer in the prefusion conformation. The RBD of each S protomer can switch between a receptor-accessible conformation known as the “up-state”and a receptor-inaccessible and buried conformation that packs against the N-terminal domain (NTD) of the neighboring protomer called the “down-state”(7). Two regions can be identified in the RBD, a conserved core and a more variable region termed as the receptor-binding motif (RBM, residues 438-506). The latter region contains residues that establish direct contact with hACE2, determining S protein affinity and specificity (8).

Inter-species spillover is often observed in the coronavirus family members, a phenomenon that mainly originates from amino acid mutations in the RBD that enables S to bind ACE proteins from two different host species (9, 10). Beyond S crucial role in restricting viral host infectivity, the protein is the target of potent neutralizing antibodies (11–13) with therapeutic use and the main antigenic component of vaccines (14, 15). It is of particular interest to understand how mutations and posttranslational modifications (PTM) in RBD affect viral infectivity, generate antigenic escape variants or restrict the humoral and cellular immunity.

Asparagine (Asn) deamidation is a spontaneous and irreversible frequently observed PTM (16, 17). As a result of the substitution in the Asn side-chain of the carboxamide nitrogen atom by a hydroxyl group, a mixture of aspartic and isoaspartic acid (a beta amino acid) is generated (18), introducing a negative charge and a rearrangement of the protein backbone in the latter case. The deamidation rate, which heavily depends on the primary sequence and local structure, can be estimated using bioinformatic tools that rely on different approaches such as structural constraints, machine-learning or primary sequence and disorder predictors (19–21). The fastest deamidation rate is often observed for the asparagine-glycine (NG) dipeptide in loosely structured regions, which are generally regarded as deamidation “hotspots” for which most of the methods provide accurate prediction (22).

The S protein contains multiple deamidation sites, but only some of them bear biological relevance. To identify these relevant sites, we applied a prioritization process based on four key points: (i) *In-silico* identification of hotspots and deamidation rates using the NGOME-LITE algorithm (23), (ii) conservation of the sites among *Betacoronaviruses*, (ii) location of the sites at the ACE2 binding surface, and (iv) experimental determination of the deamidation rates at specific hotspots by mass spectrometry.

A direct consequence of deamidation events in Asn residues located at the RBM is the modification of the electrostatic potential of the receptor-binding surface due to the introduction of a negative charge from the aspartic or isoaspartic carboxylate. Consequently, as the RBD is the preferential region from the S protein for neutralizing antibodies (11–13), deamidation could also affect the efficiency of the humoral immune response. With these data we generated a model that computes the proportion of deamidated protomers in a single viral particle and predicts how these species evolve. Our findings shed light on deamidation, an aging mechanism that operates on the RBM, a critical region that determines virus infectivity and may have a profound impact on vaccine development.

## Results

### Identification of deamidation hotspots in S proteins

We initially estimate the individual deamidation (Fig. 1A) half-time (t_1/2_) using the NGOME-LITE (23) algorithm, for all asparagine residues with no glycosylation probability. We further include positional information and the conservation degree of deamidation-prone Asn to generate a deamidation profile for a set of *Betacoronavirus* S proteins (Fig. 1B). We restricted our analysis to a group of S proteins from *Betacoronavirsuses* with demonstrated affinity for the hACE2 receptor, which includes SARS-CoV-2 (Wuhan), SARS-CoV (Urbani), SARS-CoV (GZ0402), the closely related Bat SARS-like CoVs RaTG13 (24) and WIV1 (25). SI Appendix, Table S1 (Genbank accession number, references and conservation).

**Fig. 1.**
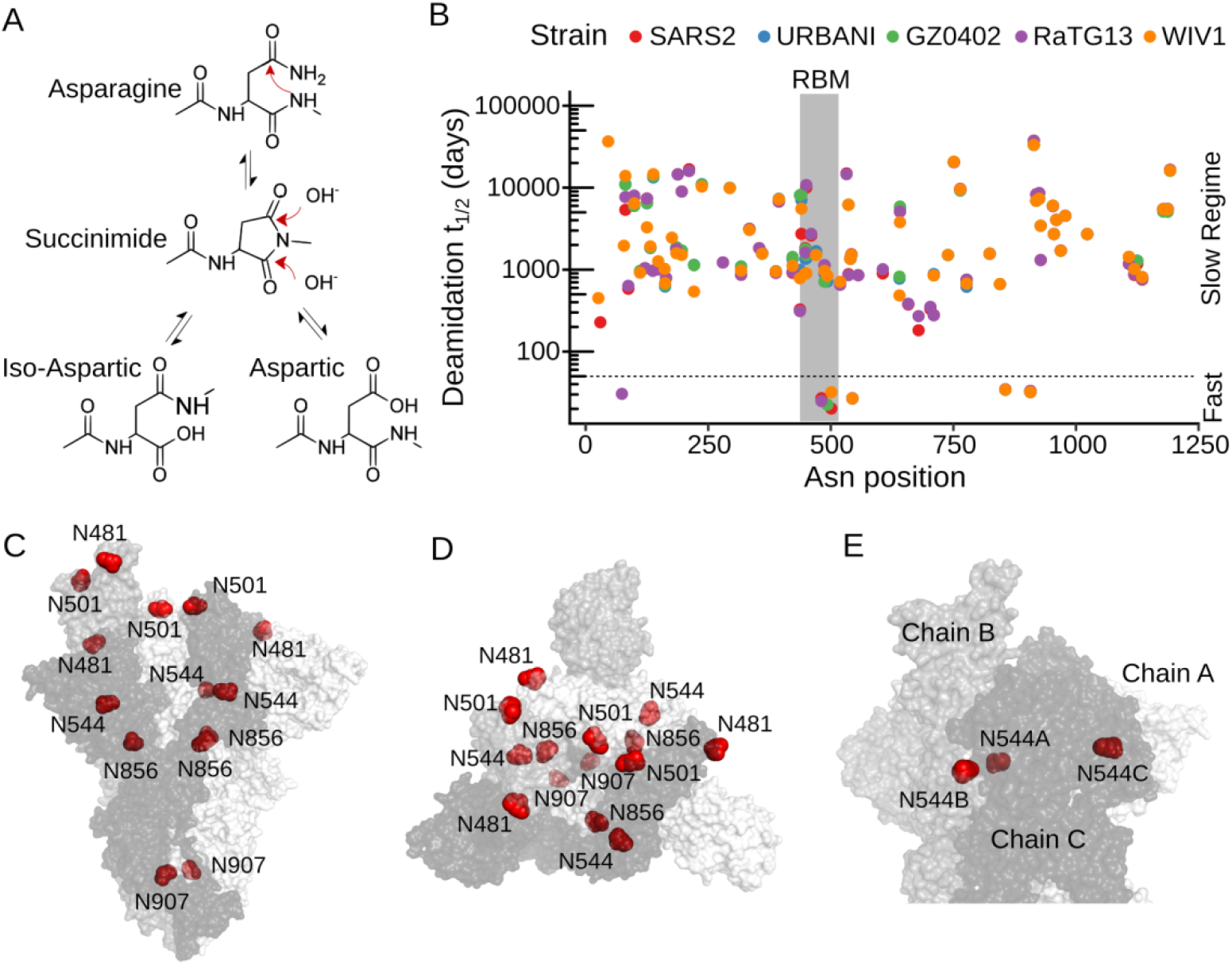
Deamidation profile of S from *Betacoronaviruses*. **A**) Schematic representation of the deamidation reaction mechanism. **B**) Deamidation profile of S proteins from a group of *Betacoronaviruses* obtained with NGOME-LITE; only non-glycosylable asparagines are included. Deamidation half-times are shown on logarithmic scale. The Asn positions are referred to the SARS-CoV-2 sequence. **C**) Deamidation hotspots on the SARS-CoV-2 S protein are shown in red. Protomers are colored in different gray tones. The PDB 6zgg (24) was used for this analysis, which contains a furin-cleaved SARS-CoV-2 S trimer, with two RBDs in the down conformation (chains A and C) and one in the up conformation (chain B). **D**) Top view of C), showing exposed deamidation hotspots at the RBD (Asn 481 and 501). **E**) Zoom of the exposed Asn 544 residues.

A similar deamidation profileis observed for the five S proteins (Fig. 1B), characterized by a major group of Asn residues in a slow-deamidation regime with estimated *t*_1/2_ greater than 100 days and a restricted group of residues in a fast-deamidation regime, with estimated *t*_1/2_ ranging between 20 to 35 days (Fig. 1B and SI Appendix, Table S2). All Asn residues in the fast-deamidation regime (hereafter referred to as deamidation hotspots) are part of NG dipeptides (SI Appendix, Fig. S1), for which a deamidation *t*_1/2_ of 1.2 days is expected in the absence of structural protection (*i.e*. in a short non-structured peptide) (18, 21) indicating that the S protein fold protects these Asn residues from deamidation. Slow regime residues virtually do not deamidate during the (expected) viral-life cycle and are devoid of any functional consequence.

Except for the single deamidation hotspot observed at position 74 in the Bat-CoV RaTG13 (in SARS-CoV-2 asparagine residue 74 is glycosylated, (26)), deamidation hotspots are restricted to a group of residues clustered between residues 481 and 501 in the RBM, including residue 493 (only observed in the GZ0402 SARS-CoV), which shows partial conservation among analyzed CoVs, and a group of strictly conserved residues at positions 544, 856 and 907 (Fig. 1C and SI Appendix, Fig. S1), observed in all the selected S sequences. Hotspots within the RBM are, altogether, termed as the *RBM deamidation cluster*. Overall, the deamidation profile shows that hotspots are not evenly distributed along these S protein sequences (Fig. 1B and Fig. 1C).

Beyond sequence context, the protein fold, backbone flexibility and surface accessibility correlate with the experimental Asn deamidation rates (27, 28). The hydrolysis of the cyclic intermediate is required to complete the deamidation reaction (Fig. 1A) for which solvent molecules must reach the reaction center. Solvent accessibility can be assessed by calculating the Relative Accessible Surface Area (RASA) of the deamidation-prone residue to a small probe radius of 1.4 Å. In addition, NGOME-LITE is unable to sense the effect of the tertiary and quaternary structure on the deamidation rate, factors that become of critical importance for hotspots centered at the inter-domain or inter-protomeric interfaces of the S trimer. Higher-order structure factors affecting deamidation rates can be identified by evaluating residue accessibility to a 3.0 Å radius probe.

To discriminate between the predicted deamidation-prone residues that are solvent accessible and those that are present at the inter-domain surface we calculated the side-chain RASA for all deamidation hotspots in the SARS-CoV-2 S structure. We performed our analysis using a structure of the furin-cleaved S protein in the pre-fusion conformation, which contains two RBDs in the down-state and one in the upstate (PDB 6zgg (24) using two probes with radii of 1.4 and 3.0 Å(29) (SI Appendix, Table S3).

All five deamidation hotspots are fully or partially accessible to the 1.4 Å radius probe (SI Appendix, Table S3), albeit to a different extent and depending on the RBD conformation, indicating that they are accessible to water molecules, a necessary requirement for hydrolysis. On the other hand, only hotspots 481 and 501 are accessible to the 3.0Å radius probe in both the up and down RBD conformations (RASA of 74.2% and 25.8% in the up conformation for Asn 481 and 501, respectively), (Fig. 1D) while Asn 544 is virtually buried in the RBD in the down-state and partially exposed (RASA of 25.6%) in the up-state (Fig. 1E).

Additionally, Asn 907 located in the heavily glycosylated stalk of S is partially accessible to the 1.4 Å radius probe through an internal channel formed by the three protomers, whereas Asn 856 is packed between the contact interface of two protomers (Fig. 1C). Both positions are fully inaccessible to the 3.0 Å radius probe, irrespective of the RBD conformation.

### Assessment of deamidation *t*_1/2_ for the 481,501 and 544 hotspots in mild conditions

The deamidation profile of SARS-CoV-2 S obtained with NGOME-LITE shows the presence of conserved deamidation hotspots at relevant protein-protein recognition interfaces. However, the fact that these sequence stretches are part of a large multidomain protein, which is not accounted for by the NGOME-LITE algorithm reduces the prediction accuracy of this bioinformatic tool.

A first experimental observation of the occurrence of deamidation at the predicted RBD hotspots was reported previously by the Wells lab (26). To monitor N-glycan occupancy in the full-length S protein, they observed 18.9% of ^18^O-Asp conversion at Asn 501, 4.8% at Asn 481 and 7.8% at Asn 544 attributable to deamidation, since these hotspots lack canonical glycosylation sequons (N-X-S/T). The amount of ^18^O-Asp conversion correlates well with the deamidation *t*_1/2_ predicted by NGOME-LITE for all five deamidation hotspot SI Appendix, Table S4.

As the deamidation kinetics is critical to evaluate the potential effect of the aspartic/isoaspartic formation in the S function, we experimentally assessed the deamidation *t*_1/2_ for the hotspots 481, 501 and 544 in a recombinantly expressed extended version of the SARS-CoV-2 RBD construct encompassing residues 319 to 566 (30) at 4 °C and 37 °C.

It should be noted that residue 544 is not part of the RBD but instead it belongs to the structural conserved subdomain 1 (SD1) that encompasses residues ~516 to ~591 not fully present in our construct. Hence, SD1 may be devoid of its native structure and the interpretation of the observed *t*_1/2_ for the 544 hotspot should be treated with caution. In our experimental set-up, the recombinant RBD protein was incubated at pH 7.4 over several days at two different temperatures and the presence of deamidated species was identified and quantified by mass spectrometry (31) (SI Appendix, Fig. S2 and Table S5). Fig. 2A shows the time-decay of the unmodified peptides (Asn containing peptides) bearing the deamidation hotspot 481, 501 and 544.

**Fig. 2.**
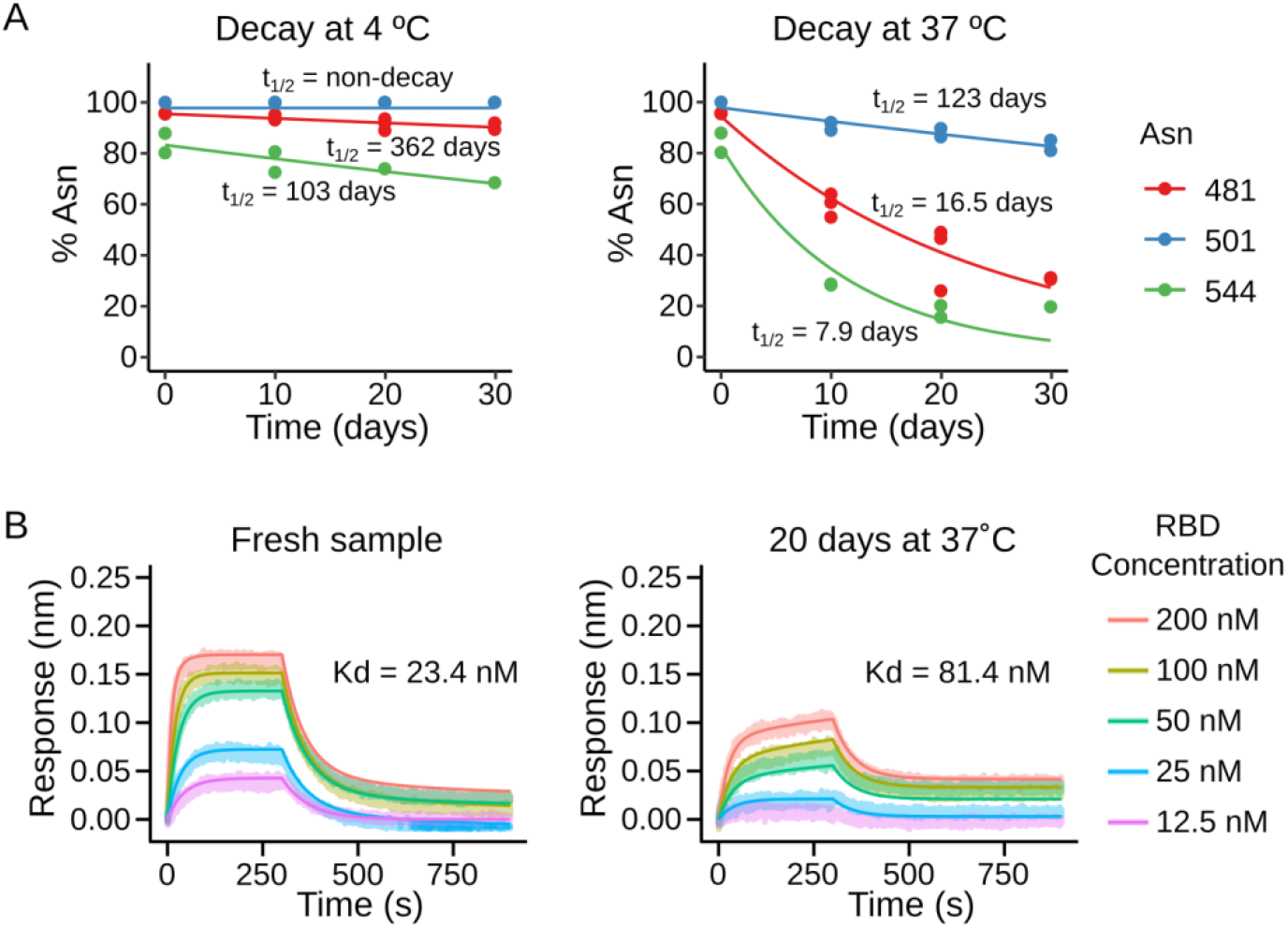
Experimental deamidation kinetic and affinity to hACE2 of an aged RBD sample. **A) Left:** Time-decay of asparagine-containing peptides for the 481, 501 and 544 hotspots at 4 °C. The lines are fit to an exponential function with initial amplitude of 100% and endpoint of 0%. **Right:** Time-decay of asparagine-containing peptides for the 481, 501 and 544 hotspot at 37 °C. The lines are fits to an exponential decay. **B) Left:** BLI response curves of a fresh RBD sample to an immobilized hACE2. **Right:** BLI response curve of an aged RBD sample (20 days at 37°C).

Initially (time = 0), the unmodified peptides covering the 481, 501 and 544 hotspots account for 95.6%, 100.0% and 84.0 % of the detected peptides, respectively (Fig. 2A and SI Appendix, Table S4); these values dropped to 34.9%, 87.9% and 17.4 % for the 481, 501 and 544 hotspots, after 20 days of incubation at 37°C. Protein conformations that are heavily influenced by ligand binding or medium conditions may explain the differences observed in deamidation propensity under different experimental conditions.

Deamidation half-times of 16.5±3.7, 123±23, and 7.9±1.2 days were obtained at 37°C for the 481,501 and 544 hotspots. Under our experimental conditions, the RBD hotspot 501 remains stable against the deamidation process despite being identified *a priori* by the NGOME-LITE as the fastest deamidation hotspot. On the contrary, Asn 544 has the highest deamidation rate, with a value close to the expected deamidation rate of a NG sequence in an unstructured peptide model (1.2 days) supporting that this residue is placed in an unstructured region. The disagreement between experimental determined deamidation rates and computational predictions for hotspots that share the common NG sequence highlight that protein structure and associated dynamicsin S can fine tune the deamidation rate.

As expected, the deamidation reaction was slowed down considerably by decreasing the temperature to 4°C (SI Appendix, Table S4). The deamidation half-time for Asn 481 at 37°C compared to 4 °C is more than 20-fold higher, highlighting that temperature can critically affect the identity of the RBM.

Deamidated species for hotspots 481 and 544 elute at two different retention times (RT) in a reverse phase (RP) chromatography experiment (SI Appendix, Fig. S2), in agreement with the reaction mechanism that proposes the generation of a mixture of aspartic or isoaspartic bearing peptides. In RP chromatography, deamidated species containing isoaspartic residues are reported to elute faster than aspartic containing peptides (18) and, based on this chromatographic behavior, the deamidation reaction (20 days at 37 °C) at hotspots 481 and 544 shows an abundance of 46.6% and 68.5% of isoaspartic containing species, respectively (SI Appendix, Table S4). On the contrary, only one peak was observed for the deamidated species at Asn 501.

We then evaluated how deamidation affects the affinity of RBD for the ectodomain of hACE2. To this end we performed a bio-layer interferometry (BLI) assay using an aged and heterogeneously deamidated RBD sample with ~65%, ~12% and ~83% of hotspots 481, 501 and 544 in its deamidated form. We observed that the unaged RBD binds hACE2 with an affinity dissociation constant (K_d_) of 23.4±0.8 nM whereas the aged RBD has a reduced affinity of only 81.4±0.8 nM (Fig. 2 B) showing that deamidation is detrimental for hACE2 binding and might impact virus infectivity.

### Topological constraints drive localization of deamidation hotspots at the RBM

Due to the critical role of the RBM residues in determining hACE2 affinity and host-specificity (32, 33), we further evaluated the conservation pattern of hotspots at the RBM deamidation cluster by extending our analysis to a group of *Betacoronaviruses* representative of the subgenus *Sarbecoviruses*(34, 35) which, in addition to SARS-CoV and SARS-CoV-2, includes numerous bat and pangolin viruses (Fig. 3A and SI Appendix, Table S6).

**Fig. 3.**
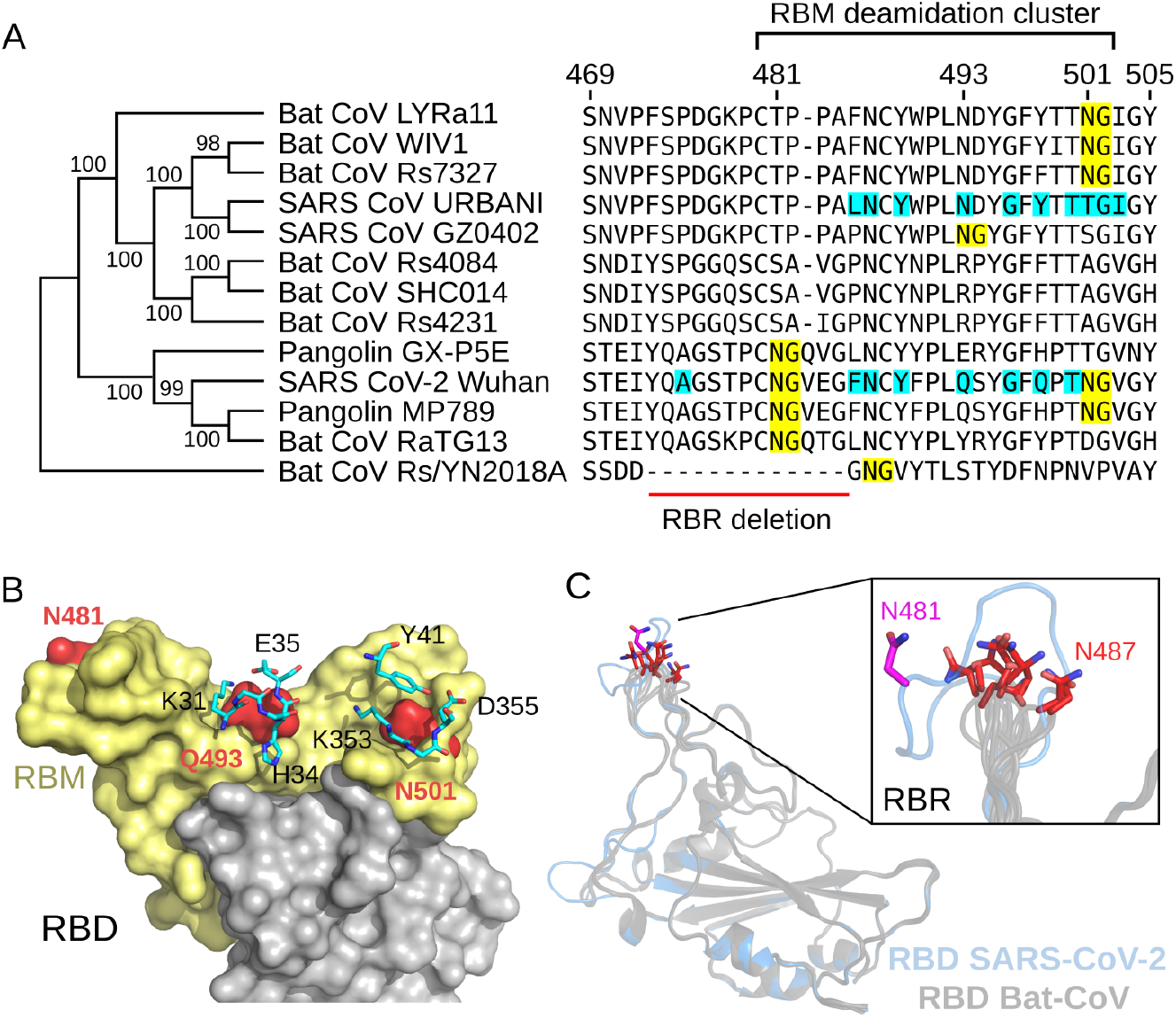
Conservation pattern of deamidation hotspots at the RBM of *Sarbecoviruses*. **A**) Cladogram of different S proteins from selected *Sarbecoronaviruses*. The RBM region is indicated and the deamidation hotspots are highlighted in yellow. Residues of SARS-CoV and SARS-CoV-2 located closer than 5Å from hACE2 residues, considered as direct contacts, are highlighted in cyan. **B**) Location of the deamidation hotspots 481, 493 and 501 at the surface of SARS-CoV-2 RBD (pdb: 6M0J). Deamidation hotspots are shown in red, RBM (residues 438-506) is shown in ocher and the core of the RBD in depicted in gray. hACE2 residues that directly interact with the RBM deamidation hotspots 493 (K31, H34 and E35) and 501 (Y41, K353 and D355) are shown as sticks. In SARS-CoV-2, the position 493 is a glutamine. **C**) Superposition of RBD from SARS-CoV-2 (pdb 6M0J, pale blue) and an ensemble of modeled structures of RBD from bat-CoV Rs/YN2018A (gray). Asn 481 residue in SARS-CoV-2 RBD is shown in magenta sticks and the Asn residues corresponding to the predicted 487 deamidation hotspot in Rs/YN2018A are drawn in red sticks. The inset details the receptor binding ridge (RBR) region.

Overall, the RBM deamidation cluster shows a variable conservation pattern (Fig. 3A) yet not random. Some *Sarbecoviruses*, including SARS-CoV Urbani and the closely related Bat CoVs Rs4084, SHC014 and Rs4231 are devoid of deamidation hotspots in their RBMs. A group including SARS-CoV GZ0402, the Bat CoVs LYRa11, WIV1, Rs7327, RaTG13, Rs/YN2018A and the pangolin GX-P5E bear one deamidation hotspot whereas SARS-CoV-2 Wuhan and Pangolin MP789 show two deamidation hotspots in their RBM.

The RBMs of the *Sarbecovirus* members can be differentiated in two groups according to the presence of a *receptor binding ridge* (RBR), which involves residues ~470 to 488 and includes a disulfide bridge (between Cys 480 and 488)(34). Altogether, deamidation hotspots in the RBMs containing a RBR are observed within an 11 amino acid stretch and are restricted to any of the following three positions: 481, 493 and 501 (Fig. 3A).

The overall folds of SARS-CoV and SARS-CoV-2 RBMs are similar with 14 residues (32) observed in equivalent positions in both structures located at less than 5Å from hACE2 binding residues (cut-off for considering a direct contact) including the 493 and 501 deamidation hotspots (Fig. 3B). Asn 493 and 501 form a direct interaction with hACE2 residues K31 and K353, respectively, which are considered critical for S binding. Mutational studies (32, 33, 36) have shown the relevance of these two positions in determining ACE2 affinity. Furthermore, and of note, the N501Y mutation that eliminates the deamidation hotspot is a hallmark of the emerging SARS-CoV-2 VOC 202012/01 (B.1.1.7) and 501Y.V2 (B.1.351) variants that are rapidly spreading in the UK and South Africa (37). Both variants are likely to have arisen independently and are associated with an increased transmissibility.

A deamidation hotspot at position 501 is repeatedly observed in SARS-CoV-2 and the pangolin CoV MP789, both sharing high sequence identity, along with the Bat CoVs LYRa11, WIV1 and Rs7327, which are more related to the SARS-CoV S protein (Fig. 3A). On the other hand, the deamidation hotspot at position 481 is fully exposed in the RBD, positioned in the external wall of the RBR (Fig. 3B and C) and does not directly participate in hACE2 binding. The deamidation hotspot 481 is also present in the closely related SARS-CoV-2, the Bat CoV RaTG13 and the two pangolin viruses CoVsGX-P5E and MP789 (Fig. 3A).

The Bat CoVRs/YN2018A, which was included as a representative *Sarbecovirus* that lacks a RBR and does not bind hACE2, shows a predicted deamidation hotspot at position 487 (SI Appendix, Table S7) that cannot be aligned with the RBR bearing sequences (Fig. 3A). The superposition of the SARS-CoV-2 RBD (pdb: 6M0J) and RBD from the Bat CoVRs/YN2018A obtained by homology modeling shows that the 481 and 487 hotspots are located in a similar position in the three-dimensional structure of the domain, suggesting that topology drives conservation of deamidation-prone asparagine residues (Fig. 3C).

### Kinetic model for S protein deamidation in the SARS-CoV-2 virion

Deamidation of the S protein RBM takes place in the context of the SARS-CoV2 life cycle. We have contextualized the results obtained for Asn 481 and 501 by numerical simulation and graphical representation of the SARS-CoV2 virion. We used the Gillespie algorithm as implemented in COPASI (38) to simulate the stepwise irreversible transition of the 33 S protein trimers in a virion (39, 40) from the intact state to the fully deamidated state. Our model considers deamidation of only Asn 481 and 501, with independent experimental half-time at 37 °C (Fig. 2). The six deamidation sites in each S trimer lead to 2^6^ possible deamidation states, which can be grouped into 20 species using symmetry considerations (SI Appendix, Fig. S3).

We performed 1000 stochastic simulations and reported the average and standard deviation of the results for each time point. Supplementary **Fig. S4** shows the full results, while Supplementary Table 8 highlights the results at several time points of interest The evolution of the different deamidated species of Spike trimers present in a virion at 37 °C are shown in Fig 4A, grouped by the total number of deamidated sites in the trimer for clarity. The intact S trimers decay with a half time of approximately four days. This value is heavily affected by temperature and the intact S trimer t_1/2_ decay at 4 °C is delayed to 110 days. Due to the multimeric nature of both the virion and the Spike protein, trimers with at least one deamidated hotspot increase early:after only one day, close to 4 S trimers would host a deamidated hotspot (mainly at Asn 481). After two days, close to 7 of the S trimers have deamidated at one site and close to an additional trimer has deamidated at two sites (SI Appendix, Table S8). Fig. 4B shows a visualization of the deamidation state of the SARS-CoV2 virion after 48 h (41). Only 17 S protein trimers are shown for clarity(41), while the population of each species is proportional to that in SI Appendix, Table S8.

**Fig. 4.**
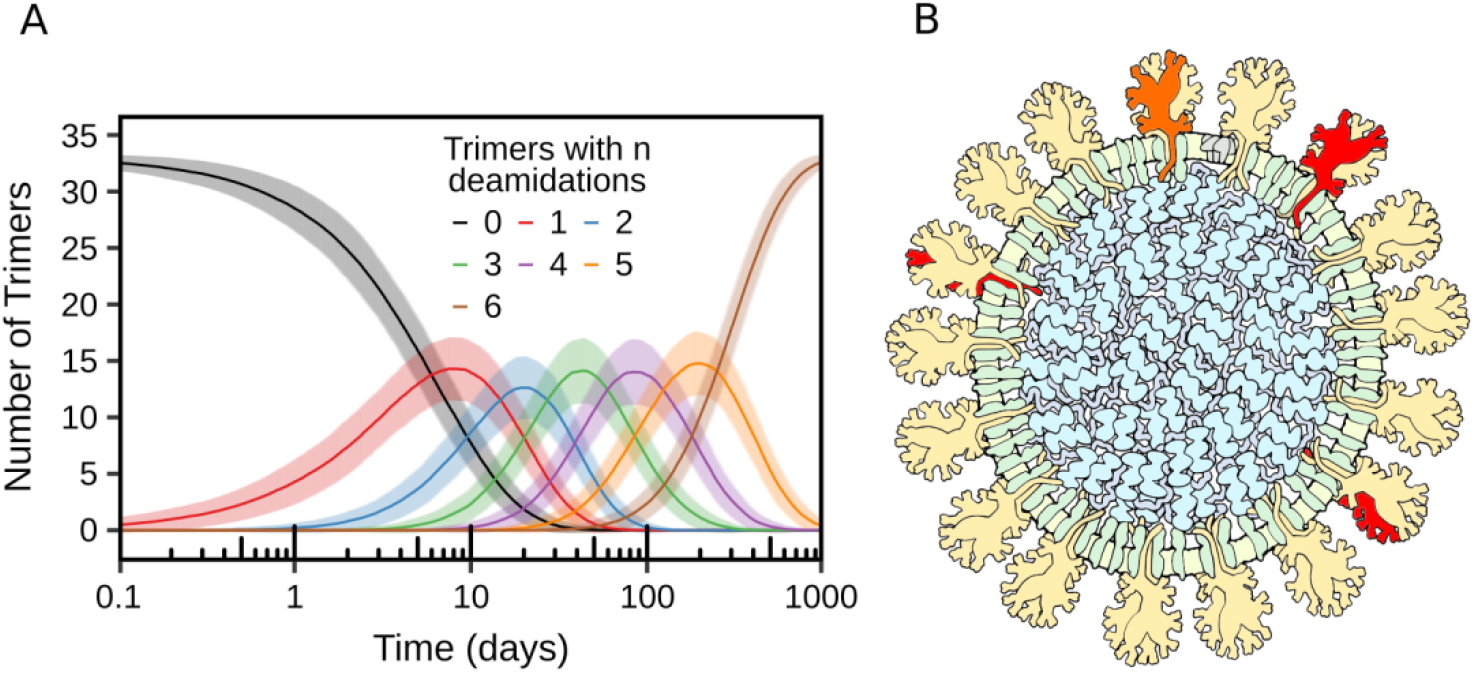
Spike protein hotspot deamidation in the context of the SARS-CoV2 virion at 37 °C. **A)** Simulated time course of the number of trimers with 0 to 6 deamidation events at sites Asn 481 and Asn 501, using the deamidation half-times from Fig 2 (see SI Appendix, Fig. S3 for simulation details). We report the average and standard deviation of 1000 simulations using the Gillespie algorithm (Supplementary SI Appendix, Fig. S4 for full results and SI Appendix, table S8 for the results at several time points of interest). **B**) Visualization of S protein deamidation in the SARS-CoV2 virion 48 h after synthesis. A cross section of the virus is shown (see (41) for details), with membrane and membranebound viral proteins M and E in green and the viral genome and associated nucleocapsid proteins N in blue. 17 of the 33 spike trimers are shown, colored according to the simulation results (SI Appendix, table S8). Yellow: undeamidated spike monomers. Red: spike monomers deamidated at Asn 481. Orange: spike monomers deamidated at Asn 501.

The intact S trimers are around four after two weeks, while the full deamidation of the virion takes about 1000 days. This timescale is far slower than that of SARS CoV-2 inactivation in solution at 37 °C, (42), indicating that deamidation of RBM hotspots is not the sole molecular mechanism for viral inactivation. This does not necessarily imply that SARS CoV-2 receptor binding is not modulated by deamidation, since a quantitative model for the effect of deamidation should include other factors such as deamidation of other sites in S, changes in RBM receptor affinity upon deamidation (Fig. 2), avidity effects arising from the trimeric nature of the S protein, modulation of the open/closed equilibrium of the trimer by deamidation, the presence of more than 30 S trimers in each virion and the presence of multiple receptor molecules on the surface of the target cell. Additionally, we would like to note that deamidation may have significantly altered the surface properties of the virion only two days after synthesis and this may have consequences for immune evasion.

## Discussion

Deamidation can critically affect protein performance and is of major importance in biologics and vaccine development, particularly when deamidating-prone asparagine residues are located in protein-protein binding interfaces or in antigenically relevant epitopes (43–45).

The deamidation profile of the S proteins from a selected group of *Betacoronaviruses* shows a discrete number of predicted deamidation hotspots that are not randomly distributed over the entire length of the S ectodomain. Instead, deamidation hotspots are observed either fully conserved or clustered within the RBM. The RBM cluster is of particular importance due to its role in hACE2 binding and includes hotspots at positions 481,493 and 501 that are partially conserved among *Sarbecovirus*. Deamidation hotspots in the RBM cluster that are critical for hACE2 binding are frequently mutated in naturally emerging variants of SARS-CoV or the SARS-CoV-2, suggesting that they are particularly sensitive to selective pressure.

We experimentally observed that Asn 481 deamidates with a half-time of 16.5 days at 37°C, a reaction that is highly temperature sensitive. An aged RBD sample, in which 65% of the 481 hotspot is observed in its deamidated form showed a 3.5-fold increase of the K_d_ for hACE2. However, the idea that deamidated residues are always detrimental for protein fitness collides with the observation that hotspot 481 is conserved among *Betacoronaviruses*. Topological constraints are likely to drive the conservation of a deamidation hotspot at position 487, resembling the 481 position of SARS CoV-2, in S protein of related bat *Betacoronaviruses* that have suffered a deletion of the entire RBR region, a critical element for hACE2 binding.

The deamidation hotspot at position 501 is in direct contact with the K353 residue in hACE2 (32) and different observations reveal its critical role in determining hACE2 affinity (46, 47).The N501Y mutation that eliminates this deamidation hotspot is a hallmark of the emerging SARS-CoV-2 variants, such as the B.1.1.7 (UK variant), the B.1.351 (South Africa variant) and the P.1 (Brazil variant) (48).

In our experimental setup we did not find any significant deamidation of Asn 501 in the SARS CoV-2 RBD construct. However, experimental evidence about deamidation of Asn 501 was reported previously for this protein (26). Such discrepancies may arise in the use of different experimental conditions that may affect protein conformation and hence, the deamidation rate. Alternatively, deamidation at these sites might not occur in the native, well folded S protein, but might be rapidly triggered when the protein is proteolyzed during the antigenic presentation process, interfering with the host-immune response (45).

Our integral kinetic model for the virion deamidation illustrates how the multimeric nature of S protein and the virion affect the accumulation of deamidate species. The model considers only two hotspots with their respective observed experimental deamidation half-times. The model shows that deamidation would not be the only molecular mechanism that reduces virus infectivity. The full deamidation time of the virion in which all of its 33 S trimers of the external surface are deamidated at both hotspots is close to 1000 days, far slower than the time required to eradicate the virus infectivity by the sole incubation at 37°C, which has been reported to be of a couple of days (42). However partial deamidated species are readily accumulated within hours and after a day close to four trimers would host a deamidated protomer, mainly at Asn 481. This supports the possibility that deamidated hotspot can affect Spike fitness. For example, deamidation supports the co-existence of chemically diverse RBM populations (i.e, with different primary sequence) within the virion, some of which may enable the virus to evade antibody recognition. In this line, the E484K mutation (that coexist with the N501Y in the Brazilian SARS CoV-2 P.1 lineage (48), located nearby to Asn 481 in the RBR, has been associated with antibody escape suggesting that mutations or modifications in the RBR region may be related to an immune evasion viral mechanism (49–51).This let us speculate that conservation of deamidation hotspots in topologically equivalent positions pursues some functionality.

One might speculate that the more infective emerging variants improve their fitness by eliminating a deamidation-prone residue that, when deamidated, decrease the affinity for the receptor. On the other hand, deamidation hotspot at position 493 (Asn 479 in the SARS-CoV sequence), which is in direct contact to K31 in hACE2 (9, 32), was only observed in SARS-CoV GZ0402, which has been isolated from a handful of individuals. Unlike, the most common SARS-CoV Urbani that possesses the dipeptide Asn-Asp at positions 479 and 480, which is not expected to be a deamidation hotspot, the SARS-CoV GZ0402 bears a Asn-Gly hotspot. Interestingly, a deamidation hotspot is generated in antibody escape mutants of SARS-CoV recovered under the selective pressure of the R80 neutralizing monoclonal antibody. The escape variants bear a D480G mutation that transforms a ND slow-deamidating Asn into a NG deamidation hotspot (52, 53). In this regard, the deamidation profile may be useful for leveraging worldwide sequence datasets aiming at prioritizing the early identification of escape variants of CoVs.

## Materials and Methods

### Deamidation estimation using NGOME-LITE

The protein sequences of five S proteins were selected to investigate the presence of deamidation hotspots (SARS-2, SARS-URBANI, SARS-GZ0402, Bat-RaTG13 and Bat-WIV1). We used NGOME-LITE (23) to predict deamidation propensity of Asn residues by using predicted t_1/2_ values. To compare sites of deamidation, the protein sequences were aligned using Clustal Omega (54) with default parameters. The residue numbering of SARS-CoV-2 S were used as reference.

### SARS CoV-2 RBD *Protein expression and purification*

The SARS CoV-2 RBD (GenBank: MN908947) was expressed and purified as reported in (30). The RBD (residues 319 - 566) was expressed by transient transfection in HEK293-F containing a secretion signal, a C-terminal Sortase motif and a non-cleavable Histidine tag. Transfected cells were incubated at 37 ° with agitation at 220 rpm and 8 % CO_2_ atmosphere. Media containing secreted protein was harvested 4 days post transfection. During this “production” time, deamidation should occur adding a cumulative amount of deamidated species at zero time. All steps of the purification were performed at 4 °C, during which deamidation rate is expected to be considerably reduced. For RBD purification, an initial IMAC step was performed on a His-Trap column (all columns by GE Healthcare) using buffers A (50 mM Tris-HCl, 0.5 M NaCl, 10 mM imidazole, pH 7.4) and B (same as buffer A but with 500 mM imidazole). An additional gel filtration step was performed on a Superdex 200 Increase 10/300 GL column equilibrated with 50 mM Tris-HCl, 150 mM NaCl, pH 7.4. The protein was concentrated to 10 mg/ml and flash frozen in liquid nitrogen and stored at −80 °C until further use.

### hACE2-*Protein expression and purification*

A gene encoding hACE2 residues 1-615 followed by a C-terminal HRV 3C cleavage site and a TwinStrepII-tag was cloned in the pXLG expression vector. HEK293F cells transfected with this construct were grown in a TubeSpin^®^ bioreactor in FreeStyle293 medium for 72h at 37°C with 8% CO2 and agitation at 180 rpm. The secreted hACE2 protein was purified from the cell culture medium using a 1 ml StrepTactin Superflow high capacity cartridge (IBA). After elution, the C-terminal TwinStrepII-tag was removed by cleavage with His6-tagged HRV 3C protease, followed by an IMAC step to remove the HRV 3C protease from the sample. Finally, the untagged hACE2 protein was injected into a Superdex200 Increase 10/300 GL size exclusion chromatography column (Cytiva) equilibrated in 50 mM Hepes pH 7.2, 150 mM NaCl and 10% glycerol. The protein was concentrated to 1.9 mg/ml, flash-frozen in liquid nitrogen and stored at −80°C.

### Chemical biotinylation of hACE2

hACE2 was chemically biotinylated using the EZ-Link NHS-PEG4-Biotin kit (Sigma). The sample was first desalted using a PD-10 gravity flow column (GE Healthcare) in 20 mM sodium phosphate buffer nd 150 mM NaCl, pH 7.4. Subsequently, the sample was chemically biotinylated for 2 h on ice, using a 20-fold molar excess of biotin over the target protein. Excess biotin was removed by running the sample through a Size Exclusion Chromatography column (Superdex 200 Increase 10/300 GL). The fractions containing hACE2 were collected and concentrated to approximately 1 mg/ml. 5% v/v glycerol was added before flash-freezing, and the samples were stored at −80 °C until further use.

### Incubation conditions for deamidation rate determination

Purified RBD sample were diluted to 1 mg/ml in buffer 50 mM Tris-HCl, 150 mM NaCl and 5% v/v glycerol, pH 7.4, filtered with a pore size of 0.22 μm for minimizing bacterial growth and added in sealed SafeSel vials that were further sealed with parafilm. Samples were incubated at 4°C and 37°C. Independent aliquots were taken at different times for deamidation quantitation.

### GluC and trypsin double digest at low pH for deamidation analysis

Digestion was done at pH 5.7 to avoid extensive deamidation during sample processing. Protein samples were diluted 1:10 in 100mM ammonium acetate (Fluka), pH 5.7, containing 10% v/v acetonitrile (Fisher Scientific). Disulfide bonds were reduced with 1 mM DTT (Sigma) and 1 mM TCEP (Invitrogen) for 30 min at 37°C. Proteins were first digested with trypsin (Promega) for 1 h at 37°C (sample to enzyme ratio 1:20) and then with GluC (Promega) (1:40 ratio) for additional 3h at 37°C. The peptides were cleaned up using an OASIS HLB μElution Plate (Waters).

### Mass spectrometry

An UltiMate 3000 RSLC nano LC system (Dionex) fitted with a trapping cartridge (μ-Precolumn C18 PepMap 100, 5μm, 300 μm i.d. x 5 mm, 100 Å) and an analytical column (nanoEase™ M/Z HSS T3 column 75 μm x 250 mm C18, 1.8 μm, 100 Å, Waters) was used, coupled directly to an Orbitrap Fusion™ Lumos™ Tribrid™ Mass Spectrometer (Thermo). Peptide samples were loaded onto the pre-column with a constant flow rate of 30 μl/min for 6 minusing 0.05% v/v trifluoroacetic acid in water. Subsequently, peptides were eluted *via* the analytical column (Solvent A: 0.1% formic acid in water) with a constant flow of 0.3 μl/min, with increasing percentage of solvent B (0.1% formic acid in acetonitrile). The peptides were introduced into the Fusion Lumos *via* a Pico-Tip Emitter 360 μm OD x 20 μm ID; 10 μm tip (New Objective) and an applied spray voltage of 2.4 kV. The capillary temperature was set to 275°C. Full mass scan (MS1) was acquired with a mass range of 375-2000 m/z in profile mode in the orbitrap with resolution of 120000. The filling time was set at a maximum of 50 ms. Data dependent acquisition (DDA) was performed with the resolution of the Orbitrap set to 30000, with a fill time of 86 ms. A normalized collision energy of 34 was applied. MS2 data were acquired in profile mode.

### Data analysis

Raw data were searched against the Uniprot human (75,069 entries, June 2020) and the SARS-CoV-2 (14 entries, June 2020) reference proteome using MaxQuant 1.6.14.0 (31). The default parameters were used with the following adjustments: Deamidation of Asn and Gln, ammonia-loss at Asn were set as variable modifications, as well as oxidation of Met and acetylation of protein N-termini. The minimum peptide length was adjusted to 6 and the maximum peptide mass to 8000 Da. For digestion Trypsin/P and GluC were enabled and the missed cleavages were set to 3. Label-free quantification was enabled, with Fast LFQ disabled. IBAQ calculations were enabled. The minimum number of unique peptides was set to 1.The Maxquant output msms.txt was imported into Skyline 20.2.0.286 (55). Precursor ion chromatograms were extracted and the values of the peak areas were exported for further calculations. MS1 spectra were inspected manually using Thermo Xcalibur Qual browser 4.2.28.14.

### Biolayer interferometry (BLI)

The binding of RBD (0 days after purification or 20 days after deamidation at 37°C) to biotinylated hACE2 was measured by biolayer interferometry (BLI) using the Octet RED96 system (FortéBio). Concentrationdependent kinetic assays were performed by loading biotinylated hACE2 on streptavidin biosensors (FortéBio), pre-equilibrated in assay buffer (PBS buffer supplemented with 0.1% (w/v) BSA and 0.02% (v/v) Tween-20) for 15 min. Prior to the association, a baseline step of 300 s was performed. Subsequently, sensors were dipped in a different well containing 200, 100, 50, 25 and 12.5 nM of RBD for 600 s followed by 900 s of dissociation time in the same buffer. All experiments were carried out at 30 °C. Data were reference-subtracted only for the association and aligned with each other with an in-house python script, using a 1:1 binding model. All figures were prepared using R and ggplot2. Two independent experiment were done.

## Abbreviations

SARS: severe acute respiratory syndrome;
CoV: coronavirus

## Acknowledgments and funding sources

We acknowledge funding from Agencia Nacional de PromocionCientifica y Tecnologica (PICT 2015-1213 to I.E.S.) and Consejo Nacional de InvestigacionesCientificas y Tecnicas (I.E.S., L.G.A., L.H.O and S.K are CONICET career investigators and J.R.L. is a member of the CONICET Support Staff for Research and Development Career). We would like to acknowledge the Sample Preparation and Characterization Facility at EMBL Hamburg.L.A.D was supported by the EMBL Interdisciplinary Postdoc Program (EIPOD) under Marie Curie COFUND actions. The funders had no role in study design, data collection and analysis, decision to publish, or preparation of the manuscript.

## Notes

### Competing Interest Statement

The authors have declared no competing interest.

