## Supplementary Information for "Deamidation drives molecular aging of the SARS-CoV-2 spike receptor-binding motif"

#### Supplementary Information Text

##### Materials and Methods

**Accessible Surface.**

The surface area was calculated using GETAREA (1) on the SARS-CoV-2 S structure (PDB code 6zgg). The accessible surface was assayed by changing the radius of the probes (1.4 and 3.0Å).

**Evolutionary analysis by Maximum Likelihood method.**

The evolutionary history was inferred by using the Maximum Likelihood method and the JTT matrix-based model (2). The tree with the highest log likelihood (-8251.69) is shown in Fig. 3 (Main text). The percentage of trees in which the associated taxa clustered together is shown next to the branches. Initial tree(s) for the heuristic search were obtained automatically by applying the Neighbor-Join and BioNJ algorithms to a matrix of pairwise distances estimated using a JTT model, and then selecting the topology with superior log likelihood value. A discrete Gamma distribution was used to model evolutionary rate differences among sites (5 categories (+G, parameter = 0.4065)). The rate variation model allowed for some sites to be evolutionarily invariable ([+I], 34.12% sites). This analysis involved 13 amino acid sequences. All positions containing gaps and missing data were eliminated (complete deletion option). There were a total of 1228 positions in the final dataset. Evolutionary analyses were conducted in MEGA X (3).

**RBM Cluster, Hot-spots at positions 481, 493 and 501.**

```
Pangolin MP789      470  YQAGSTPCNGVEGFNCYFPLQSYGFHPTNGVGYQPYRVVVLSFELLKAPATVCGPKQSTN      528
Pangolin GX-P5E     471  YQAGSTPCNGQVGLNCYYPLERYGFHPTTGVNYQPFRVVVLSFELLNGPATVCGPKLSTT      530
SARS CoV-2 Wuhan    473  YQAGSTPCNGVEGFNCYFPLQSYGFQPTNGVGYQPYRVVVLSFELLHAPATVCGPKKSTN      532
Bat CoV RaTG13      473  YQAGSKPCNGQTGLNCYYPLYRYGFYPTDGVGHQPYRVVVLSFELLNAPATVCGPKKSTN      532
Bat CoV Rs/YN2018A 459  -------------GNGVYTLSTYDFNPNVPVAYQATRVVVLSFELLNAPATVCGPKLSTQ      505
Bat CoV LYRa11      464  FSPDGKPCTP-PAFNCYWPLNDYGFYTTNGIGYQPYRVVVLSFELLNAPATVCGPKLSTD      522
Bat CoV Rs4084      461  YSPGGQSCSA-VGPNCYNPLRPYGFFTTAGVGHQPYRVVVLSFELLNAPATVCGPKLSTD      519
Bat CoV SHC014      461  YSPGGQSCSA-VGPNCYNPLRPYGFFTTAGVGHQPYRVVVLSFELLNAPATVCGPKLSTD      519
Bat CoV WIV1        461  FSPDGKPCTP-PAFNCYWPLNDYGFYITNGIGYQPYRVVVLSFELLNAPATVCGPKLSTD      519
Bat CoV Rs7327      461  FSPDGKPCTP-PAFNCYWPLNDYGFFTTNGIGYQPYRVVVLSFELLNAPATVCGPKLSTD      519
SARS CoV URBANI     460  FSPDGKPCTP-PALNCYWPLNDYGFYTTTGIGYQPYRVVVLSFELLNAPATVCGPKLSTD      518
SARS CoV GZ0402     460  FSPDGKPCTP-PAPNCYWPLNGYGFYTTSGIGYQPYRVVVLSFELLNAPATVCGPKLSTD      518
Bat CoV Rs4231      460  YSPGGQSCSA-IGPNCYNPLRPYGFFTTAGVGHQPYRVVVLSFELLNAPATVCGPKLSTD      518
```

**Hot-Spot 544**

```
Pangolin MP789      529  LVKNKCVNFNFNGLTGTGVLTESSKKFLPFQQFGRDIADTTDAVRDPQTLEILDITPCSF      588
Pangolin GX-P5E     531  LVKDKCVNFNFNGLTGTGVLTTSKKQFLPFQQFGRDISDTTDAVRDPQTLEILDITPCSF      590
SARS CoV-2 Wuhan    533  LVKNKCVNFNFNGLTGTGVLTESNKKFLPFQQFGRDIADTTDAVRDPQTLEILDITPCSF      592
Bat CoV RaTG13      533  LVKNKCVNFNFNGLTGTGVLTESNKKFLPFQQFGRDIADTTDAVRDPQTLEILDITPCSF      592
Bat CoV Rs/YN2018A 506  LVKNQCVNFNFNGLKGTGVLTDSSKRFQSFQQFGRDTSDFTDSVRDPQTLEILDITPCSF      565
Bat CoV LYRa11      523  LITNQCVNFNFNGLTGTGVLTPSLKRFQPFQQFGRDFSDFTDSVRDPKTLEVLDISPCSF      582
Bat CoV Rs4084      520  LIKNQCVNFNFNGLTGTGVLTPSSKRFQPFQQFGRDVSDFTDSVRDPKTSEILDISPCSF      579
Bat CoV SHC014      520  LIKNQCVNFNFNGLTGTGVLTPSSKRFQPFQQFGRDVSDFTDSVRDPKTSEILDISPCSF      579
Bat CoV WIV1        520  LIKNQCVNFNFNGLTGTGVLTPSSKRFQPFQQFGRDVSDFTDSVRDPKTSEILDISPCSF      579
Bat CoV Rs7327      520  LIKNQCVNFNFNGLTGTGVLTSSSKRFQPFQQFGRDVSDFTDSVRDPKTSEILDISPCSF      579
SARS CoV URBANI     519  LIKNQCVNFNFNGLTGTGVLTPSSKRFQPFQQFGRDVSDFTDSVRDPKTSEILDISPCSF      578
SARS CoV GZ0402     519  LIKNQCVNFNFNGLTGTGVLTPSSKRFQPFQQFGRDVSDFTDSVRDPKTSEILDISPCSF      578
Bat CoV Rs4231      519  LIKNQCVNFNFNGLTGTGVLTSSSKRFQPFQQFGRDVSDFTDSVRDPKTSEILDISPCSF      578
```

**Hot-Spot 856**

```
Pangolin MP789      825  FIKQYGDCLGDIAARDLICAQKFNGLTVLPPLLTDEMIAQYTSALLAGTITSGWTFGAGA      884
Pangolin GX-P5E     827  FIKQYGDCLGDIAARDLICAQKFNGLTVLPPLLTDEMIAQYTSALLAGTITSGWTFGAGA      886
SARS CoV-2 Wuhan    833  FIKQYGDCLGDIAARDLICAQKFNGLTVLPPLLTDEMIAQYTSALLAGTITSGWTFGAGA      892
Bat CoV RaTG13      829  FIKQYGDCLGDIAARDLICAQKFNGLTVLPPLLTDEMIAQYTSALLAGTITSGWTFGAGA      888
Bat CoV Rs/YN2018A 802  FMKQYGECLGDINARDLICAQKFNGLTVLPPLLTDDMIAAYTAALVSGTATAGWTFGAGA      861
Bat CoV LYRa11      819  FMKQYGECLGDISARDLICAQKFNGLTVLPPLLTDEMIAAYTAALVSGTATAGWTFGAGA      878
Bat CoV Rs4084      816  FMKQYGECLGDINARDLICAQKFNGLTVLPPLLTDDMIAAYTAALVSGTATAGWTFGAGA      875
Bat CoV SHC014      816  FMKQYGECLGDINARDLICAQKFNGLTVLPPLLTDDMIAAYTAALVSGTATAGWTFGAGA      875
Bat CoV WIV1        816  FMKQYGECLGDINARDLICAQKFNGLTVLPPLLTDDMIAAYTAALVSGTATAGWTFGAGA      875
Bat CoV Rs7327      816  FMKQYGECLGDINARDLICAQKFNGLTVLPPLLTDDMIAAYTAALVSGTATAGWTFGAGA      875
SARS CoV URBANI     815  FMKQYGECLGDINARDLICAQKFNGLTVLPPLLTDDMIAAYTAALVSGTATAGWTFGAGA      874
SARS CoV GZ0402     815  FMKQYGECLGDINARDLICAQKFNGLTVLPPLLTDDMIAAYSAALVSGTATAGWTFGAGA      874
Bat CoV Rs4231      815  FMKQYGECLGDVNARDLICAQKFNGLTVLPPLLTDDMIAAYTAALVSGTATAGWTFGAGA      874
```

Hot-Spot 907.

```
Pangolin MP789      885  ALQIPFAMQMAYRFNGIGVTQNVLYENQKLIANQFNSAIGKIQDSLSSTASALGKLQDVV      944
Pangolin GX-P5E     887  ALQIPFAMQMAYRFNGIGVTQNVLYENQKLIANQFNSAIGKIQDSLSSTASALGKLQDVV      946
SARS CoV-2 Wuhan    893  ALQIPFAMQMAYRFNGIGVTQNVLYENQKLIANQFNSAIGKIQDSLSSTASALGKLQDVV      952
Bat CoV RaTG13      889  ALQIPFAMQMAYRFNGIGVTQNVLYENQKLIANQFNSAIGKIQDSLSSTASALGKLQDVV      948
Bat CoV Rs/YN2018A 862  ALQIPFAMQMAYRFNGIGVTQNVLYENQKQIANQFNKAISQIQESLTTTSTALGKLQDVV      921
Bat CoV LYRa11      879  ALQIPFAMQMAYRFNGIGVTQNVLYENQKQIANQFNKAISQIQESLTTTSTALGKLQDVV      938
Bat CoV Rs4084      876  ALQIPFAMQMAYRFNGIGVTQNVLYENQKQIANQFNKAISQIQESLTTTSTALGKLQDVV      935
Bat CoV SHC014      876  ALQIPFAMQMAYRFNGIGVTQNVLYENQKQIANQFNKAISQIQESLTTTSTALGKLQDVV      935
Bat CoV WIV1        876  ALQIPFAMQMAYRFNGIGVTQNVLYENQKQIANQFNKAISQIQESLTTTSTALGKLQDVV      935
Bat CoV Rs7327      876  ALQIPFAMQMAYRFNGIGVTQNVLYENQKQIANQFNKAISQIQESLTTTSTALGKLQDVV      935
SARS CoV URBANI     875  ALQIPFAMQMAYRFNGIGVTQNVLYENQKQIANQFNKAISQIQESLTTTSTALGKLQDVV      934
SARS CoV GZ0402     845  ALQIPFAMQMAYRFNGIGVTQNVLYENQKQIANQFNKAISQIQESLTTTSTALGKLQDVV      934
Bat CoV Rs4231      875  ALQIPFAMQMAYRFNGIGVTQNVLYENQKQIANQFNKAISQIQESLTTTSTALGKLQDVV      934
```

**Fig. S1. Sequence alignments showing conservation of deamidation hot-spots.**

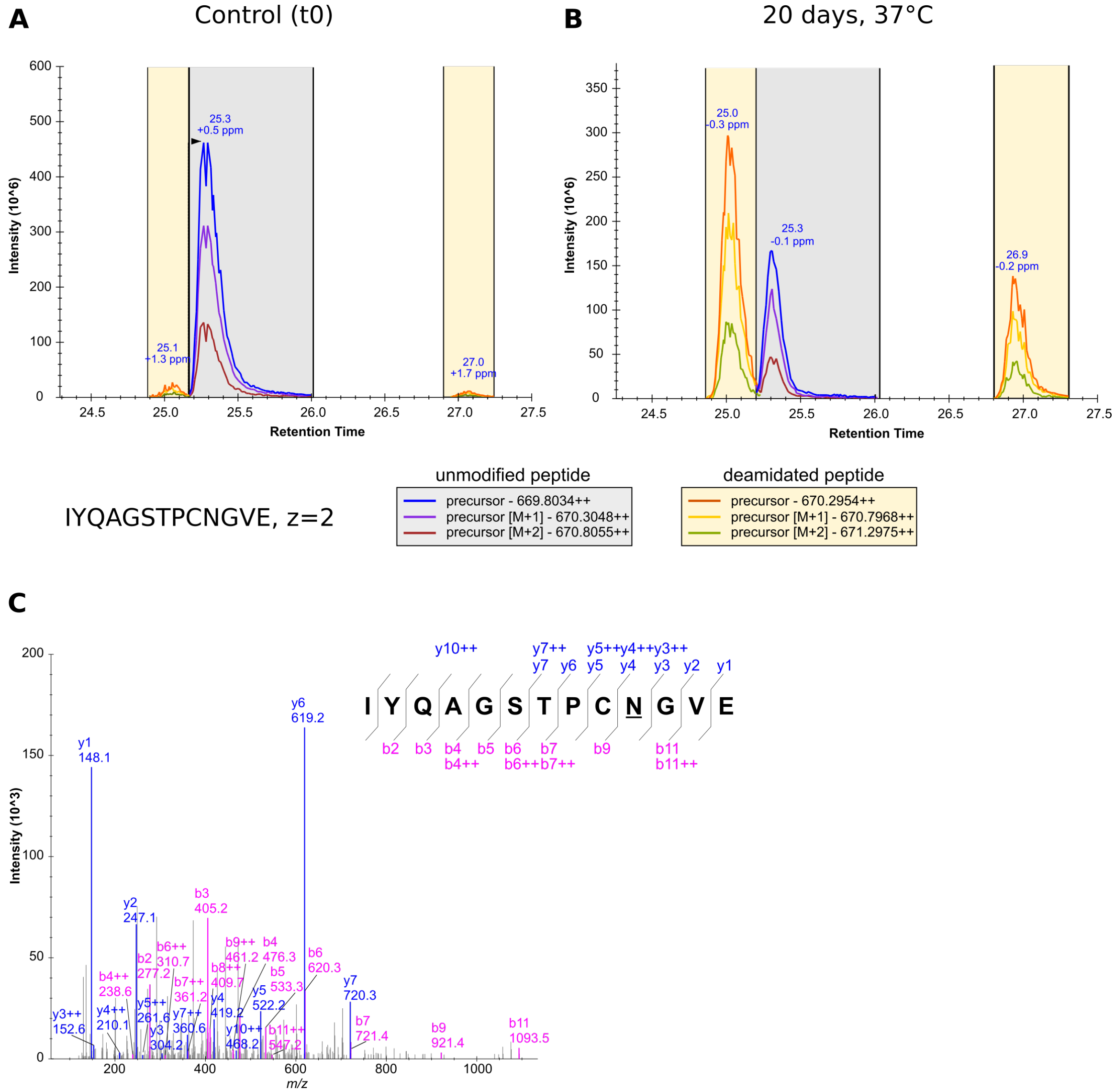


**Fig. S2**. Identification and quantification of deamidated species at the 481 hot-spot. **A**) Combined extracted ion chromatograms (XICs) of precursor ionscorresponding either to the unmodified (monoisotopic peak at 669.8034, z=2) or deamidated (monoisotopic peak at 670.2954, z=2) peptide IYQAGSTPCNGVE. At t0 only a trace of the deamidated species was observed, eluting at different retention times (RT) as compared to the unmodified peptide. **B**) Combined XIC of the deamidated and unmodified peptides obtained from an aged RBD sample. Two deamidated peptides are observed one eluting at shorter RT and the other at longer RT. **C**) The presence of an aspartic/isoaspartic acid at the position 481 was further confirmed by MS/MS.

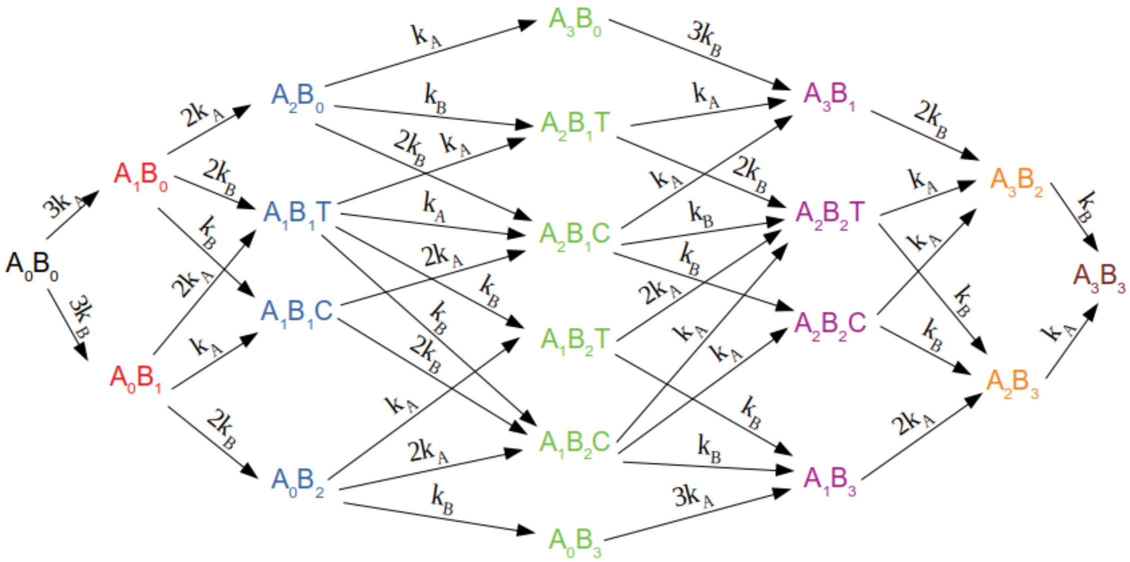


**Fig. S3. Reaction scheme for deamidation of the S protein trimer.** We considered two deamidation sites in each monomer (A and B) which denote deamidation at Asn 481 and 501, respectively. Both sites deamidate independently with microscopic reaction constants $k_A$ and $k_B$. We grouped the $2^6$ possible deamidation states into 20 species using symmetry considerations. For example, the $A_1B_1C$ species groups trimers where deamidation of one A site and one B site in the trimer took place and the two deamidation events took place in the same monomer (C stands for "cis"). The $A_1B_1T$ species groups trimers where deamidation of one A site and one B site in the trimer took place and the two deamidation events took place in different monomers (T stands for "trans"). The $A_2B_0$ species groups trimers where deamidation of two A sites and zero B sites in the trimer took place, and the $A_0B_2$ species groups trimers where deamidation of zero A sites and two B sites in the trimer took place. The 20 trimer species are colored according to the total number of deamidated sites (the color scheme is the same as in Figure 4). The reaction constants for interconversion of the 20 species take degeneracy into account. For example, the species $A_0B_0$ converts into the species $A_1B_0$ with a rate constant of $3k_A$ because $A_1B_0$ groups trimers deamidated at any of the three A sites.

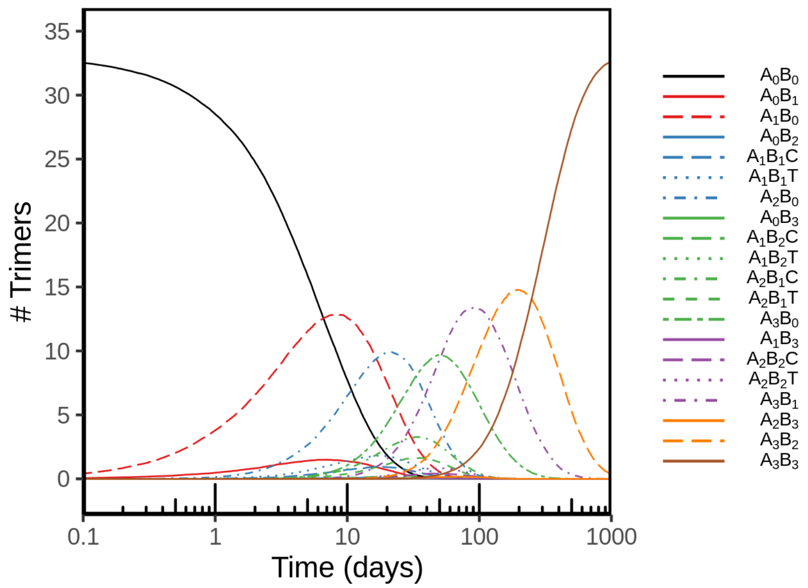


**Fig. S4. Spike protein hotspot deamidation in the context of the SARS-CoV2virion at 37 ºC.** Simulated time course of the 20 species in Fig. S3, using the deamidation half-times from Fig. 2 (see Supplementary Fig. S3 for simulation details). A and B denote deamidation at Asn 481 and 501, respectively. We report the average of 1000 simulations using the Gillespie algorithm. The color scheme is the same as in Fig. 4.

**Table S1. Betacoronaviruses with proven affinity for hACE2**.The accession numbers were obtained from GenBank (https://www.ncbi.nlm.nih.gov/genbank)

| **Virus strain** | **Accession No.** | **Host** | **Reference** |
|---|---|---|---|
| SARS-CoV-2 Wuhan-Hu-1 | YP_009724390.1 | Human<br>2019 outbreak | (4) |
| SARS-CoV Urbani | AAP13441.1 | Human<br>2002 outbreak | (5) |
| SARS-CoV GZ0402 | AY613947.1 | Human/Civet<br>2004 outbreak. No secondary infection. | (6) |
| Bat-CoV WIV1 | AGZ48831.1 | *Rhinolophus sinicus* | (7, 8) |
| Bat-CoV RaTG13 | QHR63300.2 | *Rhinolophus affinis* | (9) |

We include the SARS-CoV-2 Wuhan and SARS-CoV Urbani strains that are highly infective viruses with human-to-human transmission, the SARS-CoV GZ0402 strain isolated during the 2003/2004 episode in Guangzhou from a handful of mildly symptomatic individuals with no reported secondary transmission, alongside with human-infecting SARS coronaviruses. We include S proteins from the related Bat SARS-like CoVs RaTG13 (10) and WIV1 (8). The S proteins of SARS-CoV GZ402 share over 98% sequence identity with palm civet SARS-like CoV strains (11, 12), whereas the Bat SARS-like CoVs RaTG13 (10) and WIV1 (8) show 97.6 and 92.3% identity to the SARS CoV-2 and SARS-CoV Spike proteins, respectively.

**Table S2. Deamidation half-times ($t_{1/2}$) in days calculated from NGOME-LITE.** Positions are according to the SARS-CoV-2 numbering. Glycosylated Asn residues were excluded.

| Asn | SARS-CoV-2 Wuhan-Hu-1 | SARS-CoV Urbani | SARS-CoV GZ0402 | Bat-CoV RaTG13 | Bat-CoV WIV1 |
|---|---|---|---|---|---|
| 26 | - | - | - | - | 449.4 |
| 30 | 227.6 | - | - | - | - |
| 46 | - | - | - | - | 36650.8 |
| 74 | - | - | - | 30.5 | - |
| 78 | - | - | - | - | 1956.9 |
| 81 | 5365.3 | 11061.3 | 10851.8 | 7693.2 | 13934.6 |
| 87 | 584.5 | - | - | 634.5 | - |
| 99 | 7772.9 | 6017.4 | 5974.4 | 8012.9 | 6436.3 |
| 111 | - | 951.4 | 966.4 | - | 916.6 |
| 121 | 1042.6 | - | - | 1042.6 | - |
| 125 | 7366.1 | 6436.3 | 6515.9 | 7366.1 | 3283.1 |
| 132 | - | 1832.1 | 1858.6 | - | 1919.9 |
| 137 | 968.8 | - | - | 968.8 | - |
| 138 | - | 13302.1 | 13839.8 | - | 14607.3 |
| 148 | 1007.9 | - | - | 1007.9 | 1266.2 |
| 161 | - | - | - | - | 1021.8 |
| 162 | | 621.1 | 649.1 | - | 683.5 |
| 164 | 790.3 | - | - | 790.3 | - |
| 176 | - | - | - | - | 2458.3 |
| 185 | 1787 | 1866.8 | 1881.6 | 1779.6 | 1583.4 |
| 188 | 14594.1 | - | | 14450.2 | - |
| 196 | 8987.4 | 1707.3 | 1707.3 | 8962.9 | 1510.7 |
| 211 | 16793.8 | - | - | 15927.8 | - |
| 221 | - | 1145.2 | 1130 | - | 539.8 |
| 237 | - | 11045 | 10854.3 | - | 10298 |
| 280 | 1221.9 | - | - | 1221.9 | - |
| 294 | | 9966.9 | 9966.9 | | 9878.8 |
| 317 | 865 | 1091.5 | 1091.5 | 865 | 973.8 |
| 334 | 3152.3 | 3128.7 | 3110.6 | 3128.7 | 3044.4 |
| 354 | 1828.6 | - | - | 1824.2 | - |
| 360 | 1582.6 | 1585.8 | 1562.2 | 1577.7 | 1564.3 |
| 388 | 913.2 | 959.2 | 947.5 | 911.6 | 947.5 |
| 394 | 6749.3 | 7333.4 | 7209.2 | 6804.8 | 7209.2 |
| 422 | 914 | 1334.5 | 1430.3 | 943.5 | 1114.3 |
| 437 | 326.8 | 992.1 | 8089.9 | 311.4 | 783.5 |
| 439 | 868.1 | - | - | - | - |
| 440 | 2731 | 7035 | 8263 | - | 5538.8 |
| 448 | 1827.9 | 1352.4 | 1778 | 1616.8 | 898.5 |
| 450 | 10014.8 | 1368.2 | 1503.5 | 10778 | 898.5 |
| 460 | 2606.1 | - | - | 2731 | - |

| | | | | | |
|---|---|---|---|---|---|
| 470 | - | 1680 | 1521 | - | 1504.5 |
| 481 | 27 | - | - | 24.9 | - |
| 487 | 1125.4 | 892.7 | 711.1 | 1136.2 | 967 |
| 493 | - | 716.8 | 22.5 | - | 844.5 |
| 501 | 20.2 | - | - | - | 31.9 |
| 519 | - | 690.5 | 690.5 | 652.2 | 709.4 |
| 532 | 14947.2 | - | - | 14594.1 | - |
| 536 | 881.2 | 6191.5 | 6191.5 | 855.4 | 6191.5 |
| 540 | 1377.4 | 1353.2 | 1353.2 | 1365.2 | 1353.2 |
| 542 | 1542.3 | 1513.1 | 1513.1 | 1528.7 | 1513.1 |
| 544 | 27 | 26.8 | 26.8 | 26.8 | 26.8 |
| 556 | 855.4 | - | - | 855.4 | - |
| 606 | 897.7 | - | - | 1011 | - |
| 640 | - | 775.3 | 822.2 | - | 484.2 |
| 641 | 5103.9 | 5647 | 5883.3 | 5103.9 | 3785.2 |
| 658 | 375.5 | - | - | 384.2 | - |
| 679 | 181.9 | - | - | 271 | - |
| 703 | 330.5 | - | - | 351.8 | - |
| 710 | 280.1 | 883.6 | 849.6 | 276.9 | 849.6 |
| 739 | - | 1507.2 | 1507.2 | | 1507.2 |
| 751 | 20438.7 | 20628.1 | 20628.1 | 20438.7 | 20663.4 |
| 764 | 9655.6 | 9214.1 | 9286.8 | 9655.6 | 9537.5 |
| 777 | 754.1 | 616.9 | 685.8 | 754.1 | 675.3 |
| 824 | 1561.6 | 1571.3 | 1561.6 | 1561.6 | 1561.6 |
| 845 | - | 665.5 | 663.4 | - | 665.5 |
| 856 | 34.6 | 34.2 | 34.2 | 34.6 | 34.2 |
| 907 | 33.1 | 32 | 32 | 33.1 | 32 |
| 914 | 37409.7 | 33194.1 | 33194.1 | 37409.7 | 33194.1 |
| 919 | 8288.8 | 6906.7 | 6906.7 | 8288.8 | 6906.7 |
| 925 | 8579.8 | 7334.7 | 7334.7 | 8579.8 | 7334.7 |
| 928 | 1309.1 | 3432.1 | 3432.1 | 1309.1 | 3432.1 |
| 953 | 5953 | 6032.9 | 6032.9 | 5953 | 6032.9 |
| 955 | 2717 | 2717 | 2717 | 2717 | 2717 |
| 960 | 4047.8 | 4047.8 | 4047.8 | 4047.8 | 4047.8 |
| 969 | 1709.9 | 1701.8 | 1701.8 | 1709.9 | 1701.8 |
| 978 | 4531.3 | 4531.3 | 4531.3 | 4531.3 | 4531.3 |
| 1023 | 2713.3 | 2713.3 | 2713.3 | 2713.3 | 2713.3 |
| 1108 | 1177.9 | 1420.4 | 1420.4 | 1177.9 | 1436.5 |
| 1119 | 861 | 999.9 | 999.9 | 883.6 | 1018.2 |
| 1125 | 1173.7 | 1284.5 | 1284.5 | - | - |
| 1135 | 754.1 | 809.9 | 809.9 | 764.2 | 809.9 |
| 1178 | 5478.6 | 5478.6 | 5097.6 | 5478.6 | 5478.6 |
| 1187 | 5515.7 | 5446.4 | 5071.1 | 5515.7 | 5446.4 |
| 1192 | 16532 | 16163.5 | 16031.5 | 16532 | 16163.5 |

**Table S3. Relative Accessible Surface Area (RASA, Å$^2$) for deamidation hot-spots in SARS-CoV-2 S.**

| Asn | RASA Close Conformation Probe radius (Å) | | RASA Open conformation Probe radius (Å) | |
|---|---|---|---|---|
| | 1.4 | 3.0 | 1.4 | 3.0 |
| 481 | 74.0 | 70.1 | 90.9 | 74.2 |
| 501 | 38.3 | 25.4 | 37.9 | 25.8 |
| 544 | 15.7 | 0.0 | 40.8 | 25.6 |
| 856 | 23.4 | 2.5 | 23.8 | 2.5 |
| 907 | 30.1 | 2.0 | 34.5 | 0.5 |

Solvent Accessible Surface Area (SASA) and Relative Accessible Surface Area (RASA): SASA was calculated with GETAREA 1.0 (1) using probe radii of 1.4 and 3.0 Å for all residues in S (pdb: 6zgg, (10)) . Side chain SASA is a function of the probe radius and relative normalization was done using the SASA value at each probe radius for a fully exposed Asn residue at position 603. The SASA value for the side chain Asn 603 and a 1.4 Å probe radius is 114.74 Å$^2$, close to the 114.3 Å$^2$ reference value reported by (http://curie.utmb.edu/area_man.html) for a Asn residue in a reference tripetide Gly-Asn-Gly in a random coil conformation and calculated as an average of a 30 ensemble conformations.

**Table S4. Observed and estimated deamidation half-times for SARS-CoV-2 S hot-spots.**

| | Hot-Spot | | | | |
|---|---|---|---|---|---|
| | 481 | 501 | 544 | 856 | 907 |
| $t_{1/2}$ NGOME-LITE (days) | 27.0 | 20.2 | 27.0 | 34.6 | 33.1 |
| $^{18}$O-Asp conversion (%) # | 4.8 | 18.9 | 7.8 | 4.4 | 3.2 |
| Experimental $t_{1/2}$ (37 ºC, days) | 16.5±3.7 | 123±23 | 7.9±1.2 | n.d | n.d |
| Experimental $t_{1/2}$ (4 ºC, days) | 362±81 | No measurable decay | 103±33 | n.d | nd |
| Fold | 21.9 | - | 13.0 | - | - |
| % Unmodified species at t = 0 (4 ºC) | 95.5 | 100 | 83.2 | n.d | n.d |
| % species with a shorter RT (day 20) | 46.6 | n.o | 68.5 | n.d | n.d |
| % species with longer RT (day 20) | 18.5 | 12.0 | 14.1 | n.d | n.d |
| Sequence | NGV | NGV | NGL | NGL | NGI |

### obtained from (13)
n.d; not determined.
n.o; not observed.

**Table S5. Quantification of unmodified and deamidated species in the 481, 501 and 544 hotspots.**. Normalized total area MS1 for unmodified (Unmod) asparagine-containing species, deamidated species eluting with a shorter retention time (Deam SRT) and deamidated species eluting at larger retention times ( Deam LRT). Replicates are shown ( Exp).. n.d, not detected.

| Time (days) | Exp | Hot-spot 481 | | | Hot-spot 501 | | | Hot-spot 544 | | |
|---|---|---|---|---|---|---|---|---|---|---|
| | | IYQAGSTPCNGVE | | | GFNCYFPLQSYGFQPTNGVGYQPYR | | | CVNFNFNGLTGTGVLTE | | |
| | | t = 4 ˚C | | | | | | | | |
| | | Deam SRT | Unmod | Deam LRT | Deam SRT | Unmod | Deam LRT | Deam SRT | Unmod | Deam LRT |
| | 1 | 3.13E+08 | 1.02E+10 | 1.42E+08 | 0.00E+00 | 0.00E+00 | 0.00E+00 | 0.00E+00 | 0.00E+00 | 0.00E+00 |
| 0 | 2 | 6.13E+08 | 1.93E+10 | 2.95E+08 | 0.00E+00 | 7.37E+08 | 0.00E+00 | 1.29E+08 | 1.28E+09 | 4.96E+07 |
| | 3 | 0.00E+00 | 0.00E+00 | 0.00E+00 | 0.00E+00 | 0.00E+00 | 0.00E+00 | 3.44E+08 | 1.90E+09 | 1.28E+08 |
| | 1 | 5.07E+08 | 9.92E+09 | 2.25E+08 | 0.00E+00 | 7.26E+08 | 0.00E+00 | 0.00E+00 | 0.00E+00 | 0.00E+00 |
| 10 | 2 | 9.06E+08 | 1.89E+10 | 4.55E+08 | 0.00E+00 | 3.32E+08 | 0.00E+00 | 2.43E+08 | 1.37E+09 | 8.92E+07 |
| | 3 | 6.96E+08 | 1.92E+10 | 3.04E+08 | | | | 7.25E+08 | 2.52E+09 | 2.33E+08 |
| | 1 | 8.06E+08 | 9.47E+09 | 3.77E+08 | 0.00E+00 | 9.18E+08 | 0.00E+00 | 0.00E+00 | 0.00E+00 | 0.00E+00 |
| 20 | 2 | 1.04E+09 | 1.86E+10 | 5.91E+08 | 0.00E+00 | 2.94E+08 | 0.00E+00 | 3.69E+08 | 1.33E+09 | 1.03E+08 |
| | 3 | 8.51E+08 | 1.89E+10 | 4.54E+08 | 0.00E+00 | 0.00E+00 | 0.00E+00 | 0.00E+00 | 0.00E+00 | 0.00E+00 |
| | 1 | 0.00E+00 | 0.00E+00 | 0.00E+00 | 0.00E+00 | 8.62E+08 | 0.00E+00 | 0.00E+00 | 0.00E+00 | 0.00E+00 |
| 30 | 2 | 1.41E+09 | 1.81E+10 | 7.43E+08 | 0.00E+00 | 3.92E+08 | 0.00E+00 | 4.94E+08 | 1.31E+09 | 1.12E+08 |
| | 3 | 1.12E+09 | 1.86E+10 | 5.23E+08 | 0.00E+00 | 0.00E+00 | 0.00E+00 | 0.00E+00 | 0.00E+00 | 0.00E+00 |

| | | | | | | | | | | |
|---|---|---|---|---|---|---|---|---|---|---|
| t = 37 ˚C | | | | | | | | | | |
| 10 | 1 | 3.36E+09 | 5.82E+09 | 1.47E+09 | 0.00E+00 | 0.00E+00 | 0.00E+00 | 0.00E+00 | 0.00E+00 | 0.00E+00 |
| | 2 | 4.50E+09 | 1.29E+10 | 2.83E+09 | 0.00E+00 | 8.27E+08 | 1.03E+08 | 8.80E+08 | 4.25E+08 | 2.02E+08 |
| | 3 | 5.83E+09 | 1.22E+10 | 2.18E+09 | 0.00E+00 | 2.51E+08 | 2.19E+07 | 1.61E+09 | 7.95E+08 | 4.49E+08 |
| 20 | 1 | 5.47E+09 | 2.72E+09 | 2.46E+09 | 0.00E+00 | 0.00E+00 | 0.00E+00 | 0.00E+00 | 0.00E+00 | 0.00E+00 |
| | 2 | 1.00E+10 | 6.21E+09 | 3.96E+09 | 0.00E+00 | 7.27E+08 | 8.41E+07 | 7.12E+08 | 2.21E+08 | 1.85E+08 |
| | 3 | 7.82E+09 | 9.82E+09 | 2.56E+09 | 0.00E+00 | 3.49E+08 | 5.55E+07 | 1.68E+09 | 3.46E+08 | 2.65E+08 |
| 30 | 1 | 0.00E+00 | 0.00E+00 | 0.00E+00 | 0.00E+00 | 0.00E+00 | 0.00E+00 | 0.00E+00 | 0.00E+00 | 0.00E+00 |
| | 2 | 8.04E+09 | 9.33E+09 | 2.84E+09 | 0.00E+00 | 7.45E+08 | 1.76E+08 | 4.17E+08 | 1.39E+08 | 1.67E+08 |
| | 3 | 1.06E+10 | 6.08E+09 | 3.51E+09 | 0.00E+00 | 2.87E+08 | 5.05E+07 | 0.00E+00 | 0.00E+00 | 0.00E+00 |

**Table S6. Selected *Sarbecoronaviruses*.**

| Virus Strain | Accession | Host |
|---|---|---|
| Pangolin-CoV GX-P5E/2017 | QIA48641.1 | *Manis javanica* |
| Pangolin-CoV MP789/2020 | QIG55945.1 | *Manis javanica* |
| Bat-CoV LYRa11 | AHX37558.1 | *Rhinolophus affinis* |
| Bat-CoV Rs7327 | ATO98218.1 | *Rhinolophus sinicus* |
| Bat-CoV Rs4231 | ATO98157.1 | *Rhinolophus sinicus* |
| Bat-CoV Rs4084 | ATO98132.1 | *Rhinolophus sinicus* |
| Bat-CoV SHC014 | AGZ48806.1 | *Rhinolophus sinicus* |
| Bat-CoV Rs/YN2018A | QDF43820.1 | *Rhinolophus affinis* |

**Table S7. NGOME-LITE estimated deamidation half-time of hotspots observed in the RBM of *Sarbecoronaviruses*.**

| Virus Strain | Deamidation $t_{1/2}$ (days) | | | |
|---|---|---|---|---|
| | 481 | 487 | 493 | 501 |
| Bat-CoV Rs/YN2018A | - | 12.3 | - | - |
| Bat-CoV LYRa11 | - | - | - | 23.25 |
| Bat-CoV WIV1 | - | - | - | 31.9 |
| Bat-CoV Rs7327 | - | - | - | 23.75 |
| SARS-CoV Urbani | - | - | - | - |
| SARS-CoV GZ0402 | - | - | 22.5 | - |
| Bat-CoV Rs4084 | - | - | - | - |
| Bat-CoV SHC014 | - | - | - | - |
| Bat-CoV Rs4231 | - | - | - | - |
| Pangolin-CoV GX-P5E/2017 | 26.5 | - | - | - |
| SARS-CoV-2 Wuhan-Hu-1 | 27 | - | - | 20.2 |
| Pangolin-CoV MP789/2020 | 23.08 | - | - | 17.58 |
| Bat-CoV RaTG13 | - | - | - | 24.9 |

**Table S8. S protein hotspot deamidation in the context of the SARS-CoV2 virion at 37 ºC.** For details see Fig. S4 legend

| | Time (Days) | | | | | |
|---|---|---|---|---|---|---|
| | 0 | 1 | 2 | 3 | 7 | 14 |
| $A_0B_0$ | 33.000 ± 0.000 | 28.501 ± 2.008 | 24.789 ± 2.554 | 21.407 ± 2.662 | 11.819 ± 2.772 | 4.348 ± 1.953 |
| $A_0B_1$ | 0.000 ± 0.000 | 0.480 ± 0.696 | 0.820 ± 0.939 | 1.137 ± 1.056 | 1.498 ± 1.146 | 1.140 ± 1.034 |
| $A_0B_2$ | 0.000 ± 0.0000 | 0.004 ± 0.0632 | 0.007 ± 0.0834 | 0.017 ± 0.1293 | 0.052 ± 0.2221 | 0.085 ± 0.2790 |
| $A_0B_3$ | 0.000 ± 0.0000 | 0.000 ± 0.0000 | 0.000 ± 0.0000 | 0.000 ± 0.0000 | 0.000 ± 0.0000 | 0.001 ± 0.0316 |
| $A_1B_0$ | 0.000 ± 0.000 | 3.804 ± 1.876 | 6.555 ± 2.319 | 8.713 ± 2.457 | 12.674 ± 2.771 | 10.830 ± 2.758 |
| $A_1B_1C$ | 0.000 ± 0.000 | 0.024 ± 0.160 | 0.071 ± 0.261 | 0.135 ± 0.365 | 0.490 ± 0.690 | 0.867 ± 0.921 |
| $A_1B_1T$ | 0.000 ± 0.000 | 0.035 ± 0.184 | 0.153 ± 0.392 | 0.282 ± 0.528 | 1.004 ± 0.999 | 1.668 ± 1.256 |
| $A_1B_2C$ | 0.000 ± 0.0000 | 0.000 ± 0.0000 | 0.001 ± 0.0316 | 0.004 ± 0.0632 | 0.033 ± 0.1787 | 0.136 ± 0.3683 |
| $A_1B_2T$ | 0.000 ± 0.0000 | 0.000 ± 0.0000 | 0.003 ± 0.0547 | 0.006 ± 0.0773 | 0.021 ± 0.1435 | 0.081 ± 0.2907 |
| $A_1B_3$ | 0.000 ± 0.0000 | 0.000 ± 0.0000 | 0.000 ± 0.0000 | 0.000 ± 0.0000 | 0.000 ± 0.0000 | 0.004 ± 0.0632 |
| $A_2B0$ | 0.000 ± 0.000 | 0.146 ± 0.370 | 0.566 ± 0.734 | 1.198 ± 1.044 | 4.302 ± 1.854 | 8.586 ± 2.541 |
| $A_2B_1C$ | 0.000 ± 0.0000 | 0.003 ± 0.0547 | 0.013 ± 0.1133 | 0.031 ± 0.1791 | 0.360 ± 0.5803 | 1.479 ± 1.1743 |
| $A_2B_1T$ | 0.000 ± 0.0000 | 0.000 ± 0.0000 | 0.003 ± 0.0547 | 0.017 ± 0.1293 | 0.197 ± 0.4317 | 0.724 ± 0.8249 |
| $A_2B_2C$ | 0.000 ± 0.0000 | 0.000 ± 0.0000 | 0.000 ± 0.0000 | 0.000 ± 0.0000 | 0.006 ± 0.0773 | 0.048 ± 0.2185 |
| $A_2B_2T$ | 0.000 ± 0.000 | 0.000 ± 0.000 | 0.000 ± 0.000 | 0.000 ± 0.000 | 0.018 ± 0.133 | 0.102 ± 0.322 |
| $A_2B_3$ | 0.000 ± 0.0000 | 0.000 ± 0.0000 | 0.000 ± 0.0000 | 0.000 ± 0.0000 | 0.001 ± 0.0316 | 0.004 ± 0.0632 |
| $A_3B_0$ | 0.000 ± 0.0000 | 0.003 ± 0.0547 | 0.019 ± 0.1366 | 0.049 ± 0.2295 | 0.464 ± 0.6657 | 2.257 ± 1.4167 |
| $A_3B_1$ | 0.000 ± 0.0000 | 0.000 ± 0.0000 | 0.000 ± 0.0000 | 0.004 ± 0.0632 | 0.060 ± 0.2418 | 0.599 ± 0.7699 |
| $A_3B_2$ | 0.000 ± 0.0000 | 0.000 ± 0.0000 | 0.000 ± 0.0000 | 0.000 ± 0.0000 | 0.001 ± 0.0316 | 0.041 ± 0.2034 |
| $A_3B_3$ | 0.000 ± 0.000 | 0.000 ± 0.000 | 0.000 ± 0.000 | 0.000 ± 0.000 | 0.000 ± 0.000 | 0.000 ± 0.000 |
